# Opposing dominance within a color gene block underpins hybrid plumage signal discordance

**DOI:** 10.1101/2021.02.18.431741

**Authors:** Devin R. de Zwaan, Jacqueline Mackenzie, Else Mikklesen, Chris Wood, Silu Wang

## Abstract

The divergence of plumage color genes is increasingly recognized as important for speciation in songbirds through its influence on social signaling. However, the behavioral mechanisms underlying the eco-evolutionary feedback that acts across species boundaries is poorly understood. The hybrid zone between *Setophaga occidentalis* (SOCC) and *S. townsendi* (STOW) in the Cascade Mountain range provides a natural observatory to test the interplay between genetics, plumage signals, and territoriality in maintaining nascent species boundaries. Recently, we found that selection within a gene block underpinning color variation (ASIP-RALY) has maintained a stable and narrow hybrid zone. Here we investigated the roles of cheek darkness and flank streaking as plumage signals during simulated territorial intrusion, two melanin-based traits underpinned by ASIP-RALY that reflect opposing dominance of SOCC and STOW alleles. We found that both plumage traits act as honest signals of body size in the territorial sex (i.e., males). The opposing dominance effects of ASIP-RALY resulted in plumage signal discordance for heterozygotes, which in turn was associated with reduced territorial performance, a fitness proxy quantified by vocal and physical responses to a decoy intruder. Taken together, this study highlights a behavioral mechanism underlying selection acting on a simple genetic architecture that shapes species boundary despite gene flow.

## Introduction

Increasing evidence suggests divergence in color genes disproportionally contribute to speciation in songbirds^1–4^, yet the underlying genetic and behavioral mechanisms remain elusive. Bird plumage is often characterized by distinct patches of pigments (e.g., carotenoids, melanins) where the patch size or color intensity can relay information about an individual’s genetic or phenotypic quality, including ‘good genes’, nutritional state, immunocompetence, parental investment, or territorial dominance^5–9^. For highly territorial species, plumage traits that accurately reflect body size, aggressiveness, or combat capabilities may be considered ‘honest’ signals of territorial performance to challengers^10–12^. Since territorial disputes are energy- and time-demanding, as well as sometimes fatal, honest plumage signals may emerge and be maintained in populations to limit costly challenges^13,14^.

Divergent plumage signals can serve as barriers to gene flow^15,16^. When plumage signals established in isolated populations come into secondary contact, divergent systems may lead to signal interference and compromised territorial performance among hybrid individuals with discordant plumage traits^17–19^. In systems where territoriality is closely tied to reproductive success, incompatible plumage signals within hybrids may therefore lead to fitness loss. Plumage signal and its genetic basis are often investigated within species or populations, but rarely across species boundaries. Here we assessed multiple plumage traits within allopatric and hybrid zone populations of two closely related wood-warbler species, *Setophaga townsendi* and *Setophaga occidentalis*, to understand the potential role of plumage signals and territorial dominance in speciation.

Divergent plumage signals may be involved in speciation between *S. townsendi* (hereafter abbreviated as ‘STOW’) and *S. occidentalis* (‘SOCC’) due to frequent heterospecific male competitions^20^ within the narrow hybrid zone along the Cascade mountain range^21–23^. As migratory songbirds, males arrive prior to females in the spring and the ability to secure and maintain high-quality territories (i.e., territorial performance) is strongly linked to mate attraction and reproductive success^24–26^. Recent evidence revealed that the key genomic differentiation between allopatric parental populations resides within a color gene block, ASIP-RALY, which pleiotropically underpins variation in multiple color traits among hybrids sampled in the hybrid zone; particularly the extent of black pigmentation (melanin) and yellow (carotenoid) coloration in the crown, cheek, breast, and flank, all of which are diagnostic for the two species^23^. Spatiotemporal analysis of ASIP-RALY allele frequency revealed selection acting on this gene block^23^, although the mechanism of selection remained unclear.

One potential mechanism underlying selection against individuals heterozygous for ASIP-RALY genotypes could be reduced hybrid territorial performance through incompatible plumage signals or the breakdown of signal reliability^27,28^. Species-specific carotenoid and melanic color features may act as plumage signals in this system, as they vary in intensity within species^23,29^ and are involved in male-male competition^24,30^. Two ASIP-RALY-associated plumage traits—flank streaking and cheek darkening—demonstrate opposing dominance: the STOW allele is dominant in flank streaking, whereas the SOCC allele is dominant in cheek darkening^23^. Based on this pleiotropic opposing dominance effect of ASIP-RALY gene block, we predicted discordant cheek and flank ancestry in heterozygous individuals. If cheek darkening and flank streaking act as signals of territorial performance within species and are divergent between species, individuals with discordant plumage signals may exhibit compromised territorial performance, resulting in reduced territorial maintenance and thus reduced fitness.

We investigated the mechanism underlying selection on the ASIP-RALY gene block that may maintain or reinforce the STOW-SOCC species boundary. Since body size is a well-established predictor of territorial maintenance and fitness^11,31^, we used it as an index of individual quality and controlled for size differences between species when addressing territorial behaviour. Specifically, we assessed: 1) if cheek and flank plumage traits predict body size within species and diverge between species, 2) if pleiotropic opposing dominance of the ASIP-RALY gene block leads to discordant plumage traits in the hybrid zone, and 3) whether hybrid individuals with discordant plumage traits demonstrate reduced territorial responsiveness or aggressiveness independent of body size.

## Results

### Plumage traits as signals of male body size

Cheek darkening and flank streaking predicted male breeding quality differently among populations. Our body size index (PC1) explained 40.5% of the variation in body mass and size traits (Figure 1A). Flank streaking was predictive of male body size in both SOCC and STOW (Table 1), as the slopes (***β*_1_**) were significantly greater than 0, although the effect was stronger in STOW (Table 1). In contrast, cheek darkening predicted body size in STOW only (Table 1). Acknowleging the small sample size of STOW in this study, we validated the body size-plumage relationship in STOW with a larger sample size using museum specimens.

**Fig. 1.**
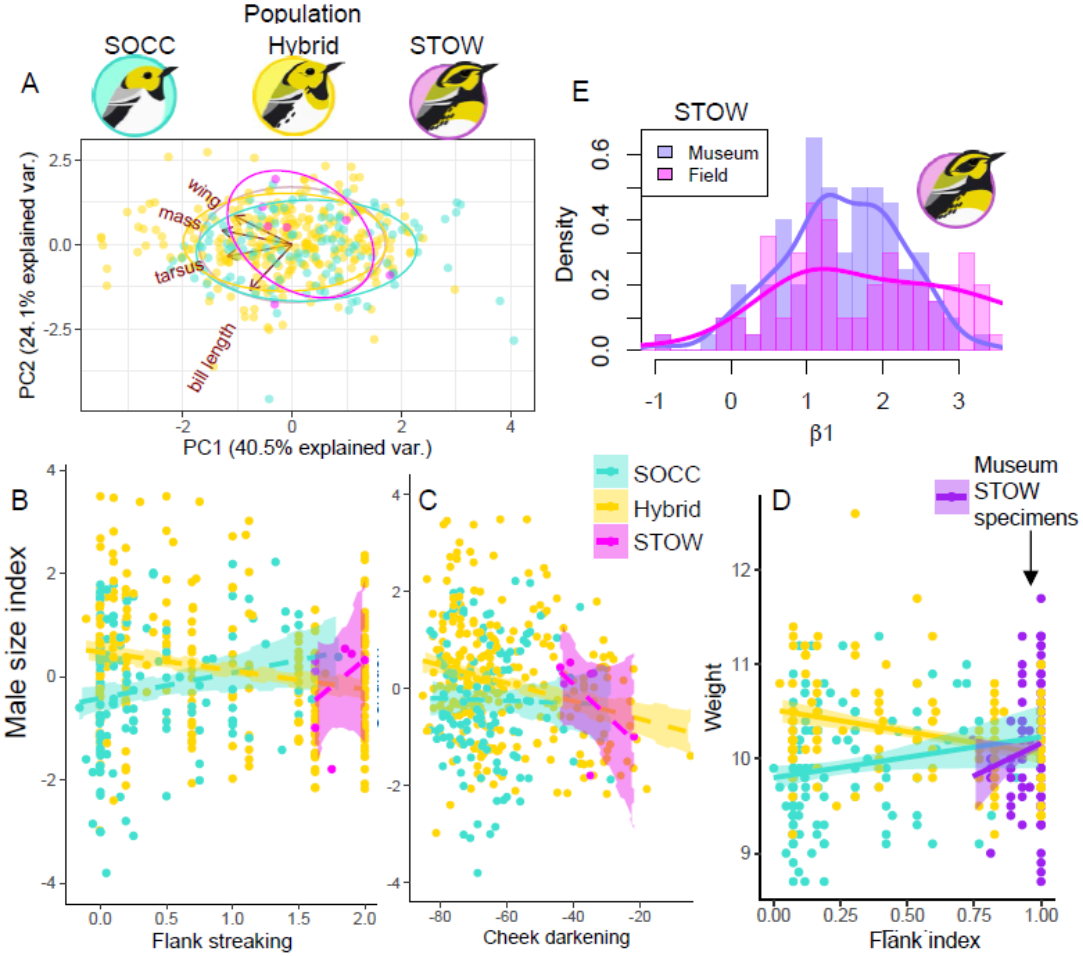
Male size signals: flank streaking and cheek coloration in the hybrid zone and parental populations (SOCC and STOW). **A**, Male size PCA, in which -PC1 was employed as a male size index. **B**, Flank streaking significantly predicts the size of males differently among populations (Table 1). **C**, Cheek darkness also predicts male size in STOW and hybrid zone (Table 1). **D**, the association of weight and flank index of hybrids, SOCC, and STOW specimens collected in British Columbia, Canada, that are stored in Burke Museum. **E**, the bootstrap distribution of slope (*β*_1_) of size-flank association in STOW (B, magenta), is similar to that of the weight-flank association in STOW museum specimens (D, purple).

**Table 1.** Bootstrap mean and 95% CI of slopes (***β*_1_**) and intercepts (***β*_0_**) of linear regression models in which the response variables were size indices and predicting variables were age-corrected cheek or flank coloration. The ***β*_1_** estimates that are significantly different from 0 are in bold.

| | Population | $\beta_0$ | $\beta_1$ | 95% CI |
| --- | --- | --- | --- | --- |
| Flank | SOCC | -0.459 | **0.523** | (0.480, 0.566) |
|  | hybrid | 0.488 | **-0.372** | (-0.392, -0.353) |
|  | STOW | -4.196 | **2.260** | (1.717, 2.803) |
| Cheek | SOCC | 0.255 | -0.017 | (-0.002, 0.002) |
|  | hybrid | 1.030 | **-0.019** | (-0.018, -0.020) |
|  | STOW | 2.555 | **-0.066** | (-0.058, -0.074) |

### Pleiotropic opposing dominance and p_discordance_

Cheek darkening and flank streaking demonstrated opposite dominance, such that the O allele was dominant for cheek darkening, while the T allele was dominant for flank streaking^23^(Figure 2A). This resulted in sickle-shaped associations between cheek darkening and flank streaking (Figure 2B) within the hybrid zone, where heterozygotes (OT) had cheek darkening similar to OO individuals, but TT-like flank streaking. As a result, we found a significant association between *p_discordance_* and ASIP-RALY genotypes among hybrids (Kruskal-Wallis rank sum test, *χ*^2^_(df=2)_ = 35.15, *p* = 2.3 × 10^-8^; Figure 2C). There was greater *p_discordance_* in heterozygotes (OT) than OO (Wilcoxon rank sum test with Bonferroni correction, *p* = 8.0 × 10^-6^) and TT (Wilcoxon rank sum test with Bonferroni correction, *p* = 3.8 × 10^-7^), while the homozygotes were not significantly different *(p* > 0.05; Figure 2C).

**Fig. 2.**
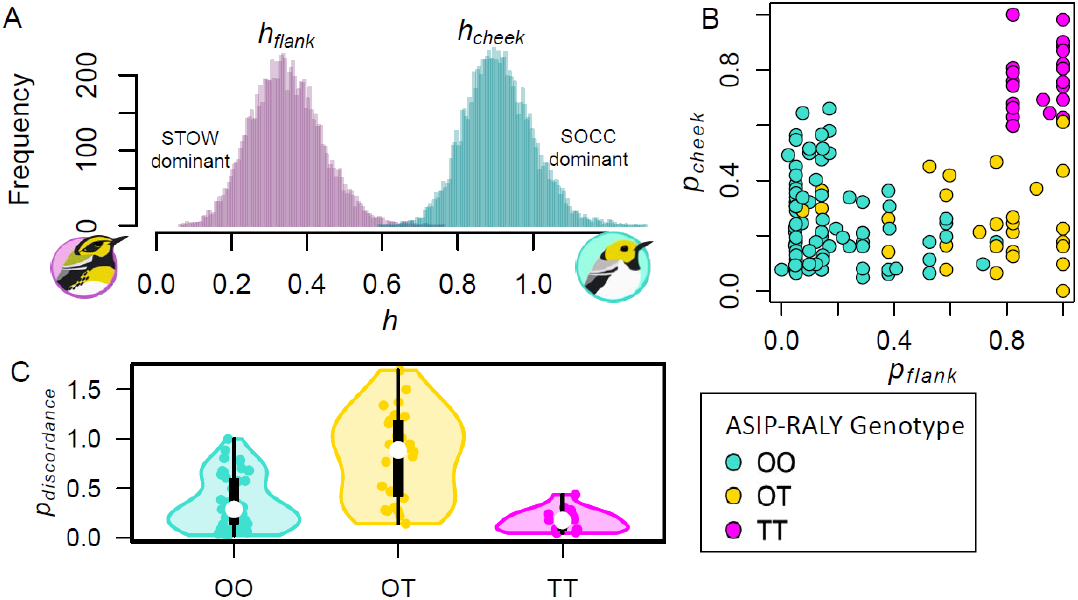
Opposing dominance in ASIP-RALY gene block leads to plumage ancestry discordance in hybrids. **A**, The distribution of dominance coefficients of flank streaking (left) and cheek darkening (right), where the STOW variant is dominant for flank streaking whereas the SOCC variant is dominant for cheek darkening^23^. **B**, Ancestry indices of cheek darkening *(p_cheek_)* and flank streaking (*p_flank_*) of individuals with ASIP-RALY genotypes: homozygous SOCC (OO), heterozygotes OT, and homozygous STOW (TT) genotype. **C**, Greater plumage ancestry discordance in individuals with OT than OO or TT.

### Plumage signal consistency

An ASIP-RALY heterozygous male tended to have a SOCC-like cheek signal but a STOW-like flank signal, which were associated with different body sizes (Table 1). Indeed, the translated signals were significantly more inconsistent in hybrids than SOCC (t = 7.8, *p* < 10^-11^) or STOW (t = 11.1, *p* < 10^-9^; Fig. 3D).

**Fig. 3.**
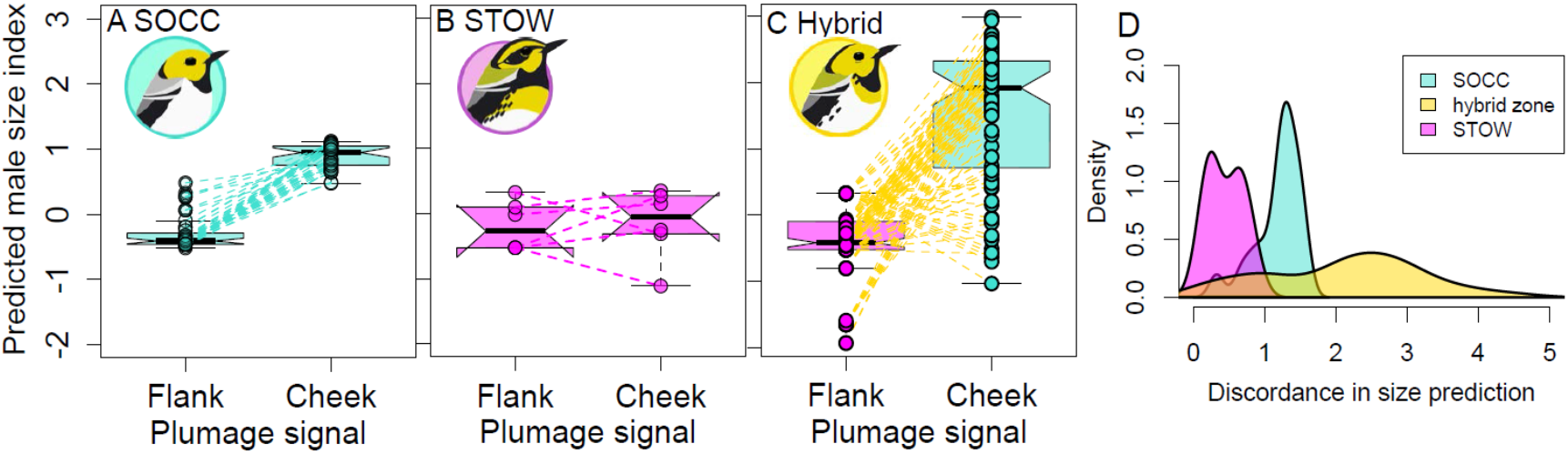
Signal information inconsistency in hybrids. **A-B**, male size prediction with cheek and flank coloration in SOCC (**A**) and STOW (**B**) parental population with a dotted line connecting the flank and cheek signal of the same male. **C**, there is hybrid signal information inconsistency relative to the parental populations (**D**, bootstrap distribution of absolute difference in size prediction based on cheek versus flank coloration).

### Territorial defense

Hybrid males with greater *p_discordance_* exhibited reduced territorial aggressiveness in response to decoy intruders. The strength of a territorial response from resident males was estimated using both physical attacks, represented by combining flight attack observations (short and longdistance flights; flight PC1), and song responses (Table S1). Flight PC1 was positively associated with greater flight attacks and explained 64.9% of the variation among males. Males with greater *p_discordance_* conducted fewer flight attacks (Spearman’s *ρ* = −0.30, *p* = 0.039; Figure 4B) and had lower song rates in response to an artificial intruder (Spearman’s *ρ* = – 0.38, *p* = 0.012; Figure 4C). Visualizing the adaptive landscape of *p_cheek_* and *p_flank_* using territorial aggressiveness as a proxy for male fitness demonstrated that males with STOW-like flanks and SOCC-like cheeks demonstrated the lowest territorial performance (Figure 4D). While there was overlap with median values of *p_cheek_* and *p_flank_*, hybrids that were more aggressive tended to have more STOW-like plumage traits (cheek and flank; Figure 4D).

**Fig. 4.**
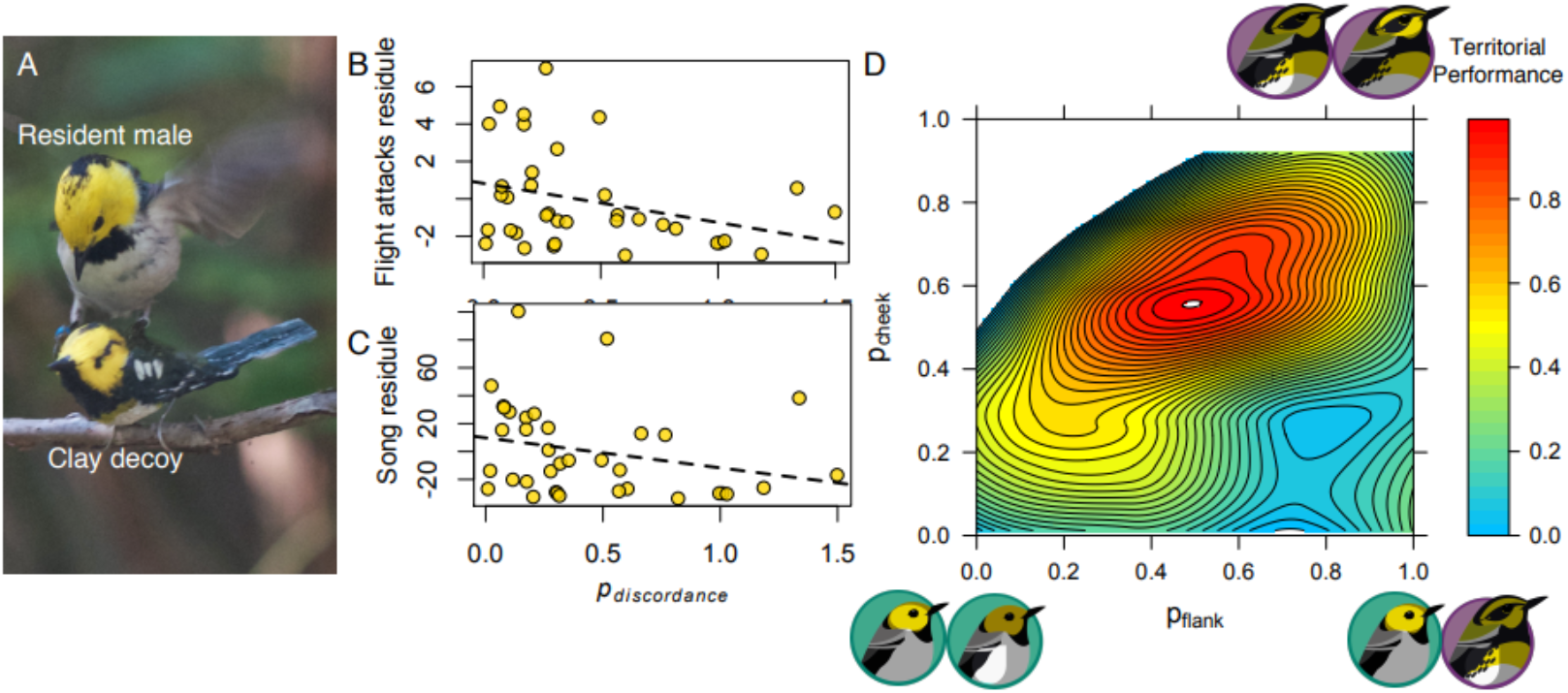
Reduced intruder territorial defense of hybrid males with greater ancestry discordance *(p_discordance_)* of plumage badges. **A**, Short flight attack of the focal male during the intruder stimulation trial (photo credit: Else Mikklesen). **B-C**, Males with greater *p_discordance_* demonstrate reduced flight attacks (B, Spearman’s *ρ* = −0.30, *p* = 0.039) and reduced song responses (C, Spearman’s *ρ* = −0.38, *p* = 0.01), independent of body size. **D**, Territorial performance with respect to the *p_flank_* and *p_cheek_* in the hybrid zone, where the males with STOW-like flanks and SOCC-like cheeks demonstrated the lowest territorial performance.

## Discussion

Here we unravel the mechanism in which the divergence of a single color gene block contributes to speciation. We found that the extent of two plumage traits, cheek darkening and flank streaking, predicts male body size in SOCC and STOW, indicating that these traits are honest signals of body condition among male conspecifics or potential mates within a social environment. Additionally, we demonstrated a pleiotropic opposing dominance effect of the ASIP-RALY gene block which led to discordant plumage signals within the hybrid zone.

Discordant plumage signals within hybrids predicted contrasting size trait information which may disrupt signal efficacy. Moreover, plumage signal discordance was associated with diminished territorial aggressiveness, represented by reduced attack and song rate in response to a decoy intruder. Taken together, these results highlight a potential behavioral mechanism contributing to selection against hybrids in a territorial context and the maintenance of the nascent species boundary between SOCC and STOW.

### Honest plumage signals

Brighter yellow cheeks and greater flank streaking were associated with larger, heavier males in both SOCC and STOW, although the strength of this relationship varied by species. Given that body size and condition traits are often associated with individual quality or social dominance^11,31,33^, this indicates that: 1) plumage traits have the potential to act as honest signals of territorial dominance in both species, but 2) the reliability of these signals may differ within each species. Evidence for plumage traits as social signals is widespread, yet the evolution and maintenance of signal honesty is likely dependent on genetic, environmental, and/or social conditions that differ among populations and species^15,34–37^.

Sexual dichromatism in carotenoid and melanin-based plumage traits, such as those present in SOCC and STOW, has been implicated in male-male competition signaling across several species^5,10,12,35,38,39^. Carotenoids are red or yellow pigments that are exogenously acquired from food or biochemically converted^8,40^. More vibrant colours therefore indicate greater resource acquisition and have been hypothesized as a signal of individual quality in males, indicating greater body condition and the ability to secure and defend resource-rich territories^6,7,38^. In contrast, melanin in black or brown patches are an expression of melanocortin systems, where melanogenesis is closely tied to intrinsic endocrinal pathways such as testosterone production^41–43^. As a result, males with darker melanistic plumage traits have been associated with greater territoriality, aggressiveness, and social dominance^44,45^.

In the context of male-male competition, honest signals that accurately reflect individual quality to territorial challengers are often referred to as ‘badges of status’^10,12^. To be considered a badge of status, plumage traits should 1) vary among counterparts of the same sex, 2) correlate with body size, condition or territorial aggressiveness of the carrier^10,13,46^, and 3) moderate aggressive interactions among counterparts by dissuading costly combats between unmatched individuals or triggering combats between individuals of similar condition^10^. Our results demonstrate support for the first two requirements: plumage traits vary among males for both species, while differences in flank streaking and cheek darkening were associated with body size and territorial aggressiveness. Manipulating the visibility of flank and cheek plumage traits and measuring how this influences the frequency of territorial challenges and success rate is required to validate the presence of badges of status in this system^11^.

### Opposing dominance of ASIP-RALY and signal discordance

Opposing dominance within the ASIP-RALY gene block^23^ resulted in discordance between cheek and flank plumage signals within hybrids, which was associated with compromised territorial performance (Figure 4). This effect is similar to the previously documented opposing dominance of two loci in *Mitoura* butterflies^47^, where the F1 hybrids (heterozygous for both loci underlying host plant preference and performance) demonstrated preference for cedar but better performance in cypress habitat. While the habitat preference and performance traits are co-adaptive within *Mitoura* species, discordance of preference and performance led to poor hybrid fitness^47^. In this *Setophaga* warbler system, opposing dominance in plumage signals occurred within one genetic region as opposed to two, resulting in signal discordance (Figure 2), information inconsistency (Figure 3), and ultimately, suboptimal territorial performance (Figure 4D). This observation is consistent with a single-locus-two-alleles model of speciation^48^, although there are other possible genomic contributors of speciation in this *Setophaga* system^49^.

Discordant signals have the potential to disrupt hybrid interactions such as male-male competition or mate attraction. Unreliable plumage signals could lead to more frequent, costly battles with aggressive, socially dominant males, potentially requiring greater investment in survival and less in the acquisition and defense of high-quality territories and mates^50,51^. Therefore, such an effect could contribute to the selection against hybrids that maintains a stable species boundary^23^. In line with this hypothesis, we found that the extent of signal discordance was associated with reduced hybrid territorial performance (Figure 4D). Territoriality is an important proxy of breeding male fitness as it is tightly associated with pairing success and return rate in SOCC and STOW males^24^. For migratory songbirds like *Setophaga* warblers, males arrive at the breeding grounds before females to acquire high-quality territories and attract mates, such that the ability to defend territories is essential to reproductive success and male fitness^25^. Our results show that hybrid males with discordant badges of status exhibited less aggressive responses to a decoy intruder, potentially indicating a reduced ability to secure and maintain territories.

This is consistent with the hybrid inferiority or parental-fitness asymmetry hypotheses, both of which posit that hybrids display inferior fitness in some way relative to their parents which in turn can maintain a narrow and stable hybrid zone^21–23,52,53^. While we assume territorial performance are associated with fitness, at least insofar as reflecting the ability to maintain territories, we could not assess breeding success or fecundity to directly link hybrid fitness with the level of signal discordance. Poor hybrid fitness may also manifest itself in other ways that we did not measure, such as reduced parental investment or survival, which could similarly limit lifetime fitness and reinforce species boundaries^54^. Importantly, body size and territorial aggression may become decoupled from reproductive fitness in the presence of alternative mating strategies (e.g., extra-pair mating) which are ubiquitous in migratory songbirds^55,56^. Future research should therefore aim to quantify hybrid fitness more directly in terms of number of offspring sired by each male, although such effort would be highly challenging as the warblers are coniferous forest canopy nesters. While additional study is needed to complete our understanding of *Setophaga* hybrid fitness, we highlight that territorial dynamics and altered social interactions in general are an intriguing mechanism by which the species boundary could be maintained.

### Aggression signals & speciation

We provide evidence for the role of male-male competition signals in speciation. Despite the long-standing view that sexual selection contributes to the evolution of reproductive isolation, signals such as plumage traits underlying male-male competition have been overlooked until recently^57–59^. The opposing dominance of ASIP-RALY leading to maladaptive discordance of hybrid badges of status is consistent with accumulating evidence that both male competition and mate choice signals are involved in reproductive isolation. Importantly, our results do not preclude that cheek and flank plumage traits may also play a role in mate choice. Recent evidence highlights the potential for some plumage traits to have dual, interacting functions, co-evolving to signal multiple receivers^35,60^. Whether this is true in the case of *Setophaga* warblers and if mate attraction may similarly be disrupted in hybrids is an important component of hybrid zone maintenance that requires further testing.

### Conclusion

We demonstrate that opposing dominance effects of ASIP-RALY led to discordant plumage signals ultimately inferior territorial performance in hybrid *Setophaga* warblers. We propose that this finding uncovers a behavioural mechanism underlying the selection on a single gene block maintains and strengthens the nascent species boundary. Additionally, our results provide valuable insights into the one-locus model of speciation in terms of how could opposing dominance within a locus/gene block result in underdominance as a simple mechanism of reproductive isolation. Such complex behavioral consequences related to a simple genetic process may represent an effective barrier to gene flow in the early stage of speciation. Future research that tracks genetic processes and social interactions over time would help reveal whether the influence of behavioral dynamics is persistent in the face of gene flow allowing speciation to proceed.

## Materials and Methods

### Study site

We carried out this study in a hybrid zone within the Cascades Mountain range of the Pacific Northwest. See Wang et al^22^ for detailed site and sampling information.

### Body size index

We sampled breeding males in allopatric populations of SOCC (N = 126) and STOW (N = 7), as well as the hybrid zone (N = 273) during the 2015 and 2016 breeding seasons. To calculate a body size index of each individual, we measured mass (g), wing length (mm), tarsus length (mm), and bill length (mm). Among the 406 males, we performed imputation to estimate missing values for body mass (n = 11), wing length (n = 1), and tarsus length (n = 2). First, we calculated a multivariate distance matrix of the four body size indices among individuals with the *dist* function in R. For each missing size variable, we adopted the value of the nearest neighbor in the multivariate distance matrix. We then used a principal component (PC) analysis (centered and scaled) with *prcomp* in R to reduce the dimensionality of the data. We selected PC1 as a measure of overall body size because it had a high factor loading (40.5%), and we took the inverse to generate the body size index where higher values represent larger and heavier males (Table S1).

**Table S1.** Loadings of each male body size variable in each PC.

|  | PC1<br>(40.5%) | PC2<br>(24.1%) | PC3<br>(19.5%) | PC4<br>(15.9 %) |
| --- | --- | --- | --- | --- |
| mass | -0.5867594 | 0.2516803 | -0.1804162 | 0.7482115 |
| tarsus | -0.5451699 | -0.2059218 | -0.6321261 | -0.5106884 |
| wing | -0.4837007 | 0.5084842 | 0.5826504 | -0.4098731 |
| bill<br>length | -0.3528978 | -0.7973062 | 0.4778965 | 0.1066817 |

**Table S2.** Loadings of each flight attack variable in the territorial intrusion trials.

|  | PC1 (64%) | PC2 (36%) |
| --- | --- | --- |
| Short flight | 0.7337579 | 0.6794110 |
| Long flight | 0.6794110 | -0.7337579 |

### Plumage coloration

We used measurements of cheek coloration and flank streaking from our previous study^23^. Briefly, for the cheek colour of breeding males, we quantified the proportion of yellow relative to black pigmentation in LAB colour space using Photoshop (Adobe Photoshop CS 2004). Mid and lower flank streaking were scored and summed to quantify total flank streaking^23^. Age correction was conducted for each of the plumage traits to control for age-related pigmentation variation. The age-correction factor was calculated as the within-quartile difference between median pigment values of each age class. We then adjusted the age-class-specific differences at each quartile of the pigment measurement with the age-correction factor^22,23^.

### Plumage signaling

To test whether cheek coloration and flank streaking predict the body size of breeding males, we fit a linear regression of the body size index against both plumage traits for each population separately (i.e., parental STOW, SOCC, or hybrid zone) to assess within-population variation:

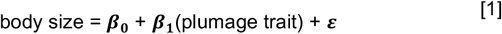

To test for a significant association between body size and plumage traits, we generated a bootstrapped mean and 95% confidence interval (CI) for β_1_ within each population by sampling individuals with replacement 100 times and re-fitting Eq. 1 for each iteration. A significant β_1_ (non-zero slope), where the 95% CI does not include zero, suggests the plumage trait is predictive of male body size.

### Museum specimens

We acknowledge that the sample size of STOW sampled in the field is small relative to the SOCC sample size (n = 126). To validate whether the associations between plumage traits and body size identified within the 7 STOW are representative of the larger parental population, we scored flank coloration of 142 existing museum specimens using the same methodology described above^21^. Museum specimens were collected in British Columbia during the breeding season (Supplementary data) and stored in the Burke Museum of Natural History and Culture (University of Washington, Seattle, Washington). Body mass for each male specimen was measured upon collection. We age-corrected the plumage scores as described above and assessed the association between mass and flank streaking using Pearson’s Product Moment correlation (r_p_). To test whether the association between flank streaking and body mass in the field samples was consistent with the museum specimens, we generated 100 bootstrapped estimates of r_p_ for both field collected STOW (n = 7) and randomly subsampled museum collected STOW (n = 7 for each iteration). If the bootstrapped distributions did not significantly differ, then the association between flank streaking and body mass for the 7 field sampled STOW was considered representative of the larger STOW population.

### Plumage color genotype

Genotype data was acquired from our previous study^23^, in which the genotypes of color gene block ASIP-RALY was denoted as OO, OT, TT for homozygous SOCC, heterozygous, and homozygous STOW, respectively. Dominance coefficients for each plumage trait were determined by the following equation:

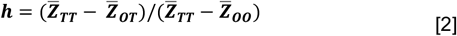

where 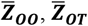, and 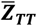 are mean phenotype individuals with OO, OT, and TT genotype within the ASIP-RALY gene block. The STOW allele is dominant if h = 0, whereas the SOCC allele is dominant if h = 1. The bootstrap distribution of h for each trait was generated by resampling individuals with replacement 10,000 times and calculating h each time. The dominance coefficients for cheek darkening and flank streaking were denoted as h_cheek_ and h_flank_, respectively.

### Plumage ancestry

Age-corrected cheek darkening and flank streaking were scaled from 0 and 1 with the following transformation function:

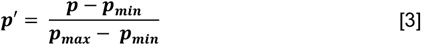

where ***p*** is the age-corrected plumage score of each individual, and ***p_min_*** and ***p_max_*** are respectively the minimum and maximum of p range. To infer hybrid ancestry, we calculated hybrid plumage ancestry (p) with the following equation:

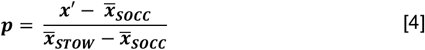

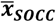 and 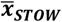 are the estimated means of plumage variation in SOCC and STOW populations. Raw ancestry scores of hybrid cheek darkening and flank streaking were scaled from 0 to 1 with Eq 3.

The ancestry discordance (p_discordance_) is the absolute difference of the scaled ancestry indices of cheek darkening (p_cheek_) and flank streaking (p_flank_):

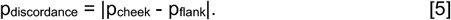

### Plumage signal consistency

To investigate signal consistency between cheek and flank plumage signals in hybrids, we separately predicted the male body size index using age-corrected cheek or flank measurements in each hybrid male based on the ***β*_0_** and ***β*_1_** estimates from Eq. 1 (Table 1). For either plumage signal, if the scaled ancestry score (p_cheek_ or p_flank_) is less than 0.5, the SOCC function was used, otherwise the STOW function was used. We then calculated plumage signal consistency as the absolute difference between body size values predicted using cheek darkening and flank streaking signals. Signal consistency was compared among hybrid and parental populations using bootstrapped distributions (100 iterations) of estimated plumage signal consistency among populations.

### Territorial performance: aggression intruder assay

To assess territorial performance of breeding hybrid males that were genotyped and had their plumage traits scored (n = 59), we quantified territorial defense of the resident males in terms of vocal (song) and physical (flight attacks) responses during an artificial territorial intrusion. We revisited territories 5—10 days after banding to carry out the territorial intrusion trials. Two researchers, one observer and one taking notes, stood 15 m from the decoy (see below). An initial quiet period of 3 min was performed to control for disturbance around the area when setting up the experiment, and then a playback consisting of a series of song recordings were played during the following 12 minutes. The response of focal males was recorded using three variables: (1) song rate, (2) short-distance flight attacks (flight around the decoy within 5 meters), and (3) long-distance flight attacks (flight around the decoy between 5 to 15 meters). Clay decoys were painted as mature individuals (> 2 years) in breeding plumage (Figure 4A) of STOW, SOCC, and hybrid (with hybrid index = 0.5) following Rowher & Wood (1998). To simulate territorial intrusion in the hybrid zone, where parental and hybrid individuals co-occur, a decoy was randomly selected as visual stimulus for each territorial intrusion trial. Decoys were presented on a tree branch that was about two meters above the ground, within 5 meters of vegetation on which the responding male could alight without being forced to contact the mount. Six individual songs were recorded with Marantz PMD660 and an Audio-Technica 815a Shotgun microphone from different populations, consisting of two songs from each of the S. townsendi allopatric populations, the hybrid zone, and the allopatric S. occidentalis zone. Playback during the territorial intrusion involved playing all six songs back-to-back in a randomly selected order (12 min in total) using a small speaker (Jam^®^ HX-P250) placed 1 meter below the mount. To consistently score territorial behaviors, a single observer (SW) described all behavioral trials.

We analyzed territorial response in terms of vocal (total song frequency) and physical response (flight attacks). To represent flight attacks, we took the PC1 in a principal component analysis with short and long flight frequencies (please see above). We then tested the association between territorial response and p_discordanc_, with body size as a control for differences between species. To account for body size as the potential confounding variable, we took the residuals of the linear regression model: territoriality = ***β*_0_** + ***β*_1_** (body size) + ***ε***. Then we tested the association between the residuals and p_discordance_ with a one-tailed Spearman’s correlation test. We predicted that individuals with greater p_discordance_ should demonstrate poor territorial performance, thus pdiscordance should be negatively correlated with territorial response.

To visualize the adaptive landscape with respect to hybrid p_cheek_ and p_flank_, we approximated male fitness using territorial aggressiveness, or the averaged flight and song response, as a proxy. To ensure that flight attacks and song responses contributed to territorial performance equally, we scaled the two variables with Eq. 3 before averaging. The bottom right corner of signal discordance corresponds to the valley or lowest territorial performance, while the ridge of performance occurs along the diagonal space of signal concordance. All analyses were conducted in R version 3.6.3 (R Core Team 2017).

## Acknowledgments

We thank Ruth Midgley for wonderful assistance in the field. We thank Darren Irwin, Sievert Rohwer, Chris Wood, Graham Coop, Sally Otto, Dolph Schluter, Loren Rieseberg, Maria Servedio, and Dahong Chen for helpful discussions. Funding was provided by NSERC to S. Wang, and Werner and Hildegard Hesse Research Awards to S. Wang and J. Mackenzie. All data were collected in accordance with UBC animal care.

## Notes

https://doi.org/10.5061/dryad.j6q573nd2

### Competing Interest Statement

The authors have declared no competing interest.

### Summary of Updates

We have included more STOW samples from the museum. In addition, we tested explicitly the effect of different intersects and slopes on size information inconsistency within each hybrid. We have also controlled for body size when estimating territorial performance of hybrids.

## References

1. Baiz, M. D., Wood, A. W., Brelsford, A., Lovette, I. J. & Toews, D. P. L. Pigmentation Genes Show Evidence of Repeated Divergence and Multiple Bouts of Introgression in Setophaga Warblers. Curr. Biol. 31, 1–7 (2021).

2. Toews, D. P. L. et al. Plumage Genes and Little Else Distinguish the Genomes of Hybridizing Warblers. Curr. Biol. 26, 2313–2318 (2016).

3. Uy, J. A. C., Moyle, R. G., Filardi, C. E. & Cheviron, Z. A. Difference in Plumage Color Used in Species Recognition between Incipient Species Is Linked to a Single Amino Acid Substitution in the Melanocortin-1 Receptor. Am. Nat. 174, 244–254 (2009).

4. Campagna, L. et al. Repeated divergent selection on pigmentation genes in a rapid finch radiation. Sci. Adv. 3, e1602404 (2017).

5. Searcy, W. A. & Nowicki, S. The evolution of animal communication: Reliability and deception in signaling systems. The Evolution of Animal Communication: Reliability and Deception in Signaling Systems (2010). doi:10.1650/0010-5422(2006)108[989:br]2.0.co;2

6. Hill, G. E. Female mate choice for ornamental coloration. Bird Color. Vol. II. Funct. Evol. (2006).

7. McGraw, K. J. & Ardia, D. R. Carotenoids, Immunocompetence, and the Information Content of Sexual Colors: An Experimental Test. Am. Nat. 162, 704–712 (2003).

8. Svensson, P. A. & Wong, B. B. M. Carotenoid-based signals in behavioural ecology: A review. Behaviour (2011). doi:10.1163/000579510X548673

9. Hill, G. E. Plumage coloration is a sexually selected indicator of male quality. Nature 350, 337–339 (1991).

10. Rohwer, S. The social significance of avian winter plumage variability. Evolution (N. Y). 29, 593–610 (1975).

11. Hagelin, J. C. The kinds of traits involved in male—male competition: a comparison of plumage, behavior, and body size in quail. Behav. Ecol. 13, 32–41 (2002).

12. Studd, V. M. & Robertson, J. R. Evidence for reliable badges of status in territorial yellow warblers (Dendroica petechia). Anim. Behav. 33, 1102–1113 (1985).

13. Rohwer, S. The Evolution of Reliable and Unreliable Badges of Fighting Ability. Am. Zool. 22, 531–546 (1982).

14. Goymann, W. & Wingfield, J. C. Allostatic load, social status and stress hormones: The costs of social status matter. Anim. Behav. 67, 591–602 (2004).

15. Hund, A. K. et al. Divergent sexual signals reflect costs of local parasites*. Evolution (N. Y). (2020). doi:10.1111/evo.13994

16. Turbek, S. P. et al. Rapid speciation via the evolution of pre-mating isolation in the Iberá Seedeater. 0256, (2021).

17. Schultz, J. K. & Switzer, P. V. Pursuit of heterospecific targets by territorial amberwing Dragonflies (Perithemis tenera Say): A case of mistaken identity. J. Insect Behav. 14, 607–620 (2001).

18. Birch, L. C. The Meanings of Competition. Am. Nat. 91, 5–18 (1957).

19. Gröning, J. & Hochkirch, A. Reproductive Interference Between Animal Species. Q. Rev. Biol. 83, 257–282 (2008).

20. Pearson, S. F. & Rohwer, S. Asymmetries in male aggression across an avian hybrid zone. Behav. Ecol. 11, 93–101 (2000).

21. Rohwer, S. & Wood, C. Three Hybrid Zones Between Hermit and Townsend ‘ S Warblers in Washington and Oregon. Auk 115, 284–310 (1998).

22. Wang, S., Rohwer, S., Delmore, K. E. & Irwin, D. E. Cross-decades stability of an avian hybrid zone. J. Evol. Biol. 32, 1242–1251. (2019).

23. Wang, S. et al. Selection on a small genomic region underpins differentiation in multiple color traits between two warbler species. Evol. Lett. 4–6, 502–515 (2020).

24. Pearson, S. F. Behavioral asymmetries in a moving hybrid zone. Behav. Ecol. 11, 84–92 (2000).

25. Morbey, Y. E. & Ydenberg, R. C. Protandrous arrival timing to breeding areas: A review. Ecol. Lett. 4, 663–673 (2001).

26. Reudink, M. et al. Non-breeding season events influence sexual selection in a long-distance migratory bird. Proc. R. Soc. B Biol. Sci. 276, 1619–1626 (2009).

27. Vortman, Y., Lotem, A., Dor, R., Lovette, I. J. & Safran, R. J. The sexual signals of the East-Mediterranean barn swallow: A different swallow tale. Behav. Ecol. 22, 1344–1352 (2011).

28. Vortman, Y., Lotem, A., Dor, R., Lovette, I. & Safran, R. J. Multiple sexual signals and behavioral reproductive isolation in a diverging population. Am. Nat. 182, (2013).

29. Jackson, W. M., Wood, C. S. & Rohwer, S. Age-Specific Plumage Characters and Annual Molt Schedules of Hermit Warblers and Townsend’s Warblers. Condor 94, 490–501 (1992).

30. Pearson, S. F., Rohwer, S., Museum, B. & Museum, B. Asymmetries in male aggression across an avian hybrid zone. Behav. Ecol. 11, 93–101 (2000).

31. Kelly, C. D. The interrelationships between resource-holding potential, resource-value and reproductive success in territorial males: How much variation can we explain? 62, 855e871 (2008).

32. R Core Team (2017). R: A language and environment for statistical computing. R Found. Stat. Comput. Vienna, Austria. R Foundation for Statistical Computing (2017). doi:/S0103-64402004000300015

33. Alatalo, R. V. & Moreno, J. Body size, interspecific interactions, and use of foraging sites in tits (Paridae). Ecology (1987). doi:10.2307/1939868

34. Safran, R. J., Adelman, J. S., McGraw, K. J. & Hau, M. Sexual signal exaggeration affects physiological state in male barn swallows. Curr. Biol. 18, R461–462 (2008).

35. Senar, J. C. Color displays as intrasexual signals of aggression and dominance. in Bird coloration (eds. Hill, G. & McGraw, K.) 87–136 (Harvard University Press, 2006).

36. McGraw, K. J. An update on the honesty of melanin-based color signals in birds. Pigment Cell and Melanoma Research 21, 133–138 (2008).

37. Tibbetts, E. A. & Safran, R. J. Co-evolution of plumage characteristics and winter sociality in New and Old World sparrows. J. Evol. Biol. 22, 2376–2386 (2009).

38. Blount, J. D. & McGraw, K. J. Signal Functions of Carotenoid Colouration. in Carotenoids 213–236 (Birkhäuser Basel., 2008). doi:10.1007/978-3-7643-7499-0_11

39. Chaine, A. S., Shizuka, D., Block, T. A., Zhang, L. & Lyon, B. E. Manipulating badges of status only fools strangers. Ecol. Lett. 21, 1477–1485 (2018).

40. Hill, G. E. Energetic constraints on expression of carotenoid-based plumage coloration. J. Avian Biol. 31, (2000).

41. McGraw, K. J., Mackillop, E. a, Dale, J. & Hauber, M. E. Different colors reveal different information: how nutritional stress affects the expression of melanin- and structurally based ornamental plumage. J. Exp. Biol. 205, 3747–3755 (2002).

42. Bókony, V., Garamszegi, L. Z., Hirschenhauser, K. & Liker, A. Testosterone and melanin-based black plumage coloration: A comparative study. Behav. Ecol. Sociobiol. 62, 1229–1238 (2008).

43. Galván, I. & Solano, F. Bird integumentary melanins: Biosynthesis, forms, function and evolution. Int. J. Mol. Sci. 17, 520 (2016).

44. Roulin, A. Condition-dependence, pleiotropy and the handicap principle of sexual selection in melanin-based colouration. Biol. Rev. 91, 328–348 (2016).

45. San-Jose, L. M. & Roulin, A. Toward understanding the repeated occurrence of associations between melanin-based coloration and multiple phenotypes. Am. Nat. (2018). doi:10.1086/698010

46. Rohwer, S. & Rohwer, F. C. Status signalling in harris sparrows: Experimental deceptions achieved. Anim. Behav. 26, (1978).

47. Forister, M. L. Independent inheritance of preference and performance in hybrids between host races of Mitoura butterflies (Lepidoptera: Lycaenidae). Evolution (N. Y). 59, 1149–1155 (2005).

48. Moore, W. S. A single locus mass-action model of assortative mating, with comments on the process of speciation. Heredity (Edinb). 42, 173–186 (1979).

49. Wang, S. et al. Signatures of mito-nuclear climate adaptation in a warbler species complex. bioRxiv (2020). doi:10.1101/2020.04.06.028506

50. Qvarnstrom, A. Experimentally increased badge size increases male competition and reduces male parental care in the collared flycatcher. Proc. R. Soc. B Biol. Sci. 264, 1225–1231 (1997).

51. Møller, A. P. Variation in badge size in male house sparrows Passer domesticus: evidence for status signalling. Anim. Behav. 35, 1637–1644 (1987).

52. Barton, N. H. & Hewitt, G. M. Analysis of Hybrid Zones. Annu. Rev. Ecol. Syst. 16, 113–148 (1985).

53. Barton, N. H. & Hewitt, G. M. Adaptation, speciation and hybrid zones. Nature (1989). doi:10.1038/341497a0

54. Servedio, M. R. & Noor, M. A. The role of reinforcement in speciation: Theory and data. Annu. Rev. Ecol. Syst. 34, 339–364 (2003).

55. Stutchbury, B. J. & Morton, E. S. The effect of breeding synchrony on extra-pair mating systems in songbirds. Behaviou. Behaviour 132, 675–690 (1995).

56. Coppack, T., Tøttrup, A. P. & Spottiswoode, C. Degree of protandry reflects level of extrapair paternity in migratory songbirds. J. Ornithol. 147, 260–265 (2006).

57. Tinghitella, R. M. et al. On the role of male competition in speciation: A review and research agenda. Behav. Ecol. 29, 783–797 (2018).

58. Pfennig, D. W. & Pfennig, K. S. Character displacement and the origins of diversity. Am. Nat. 176, S26–S44 (2010).

59. Polechová, J. & Barton, N. H. Speciation through competition: A critical review. Evolution (N. Y). 59, 1194–1210 (2005).

60. Tarof, S. A., Dunn, P. O. & Whittingham, L. A. Dual functions of a melanin-based ornament in the common yellowthroat. Proc. R. Soc. B Biol. Sci. 272, 1121–1127 (2005).

